# Pup Exposure Is Sufficient for Activating Pup Retrieval Behavior in Pup-Naive Virgin Mice

**DOI:** 10.1101/2025.11.05.686743

**Authors:** Eduarda Streit Morsch, Elizabeth A. Amadei, Roman Boehringer, Benjamin F. Grewe, Pau Vilimelis Aceituno

## Abstract

Maternal behaviors in mammals depend on complex interactions between innate neural circuits, hormonal states, and external stimuli, such as exposure and experience with offspring. Literature has shown that exposure to offspring is enough to trigger maternal behavior in alloparental (non-parental) subjects. A recent study has posited a secondary path to acquire maternal behavior through socially transmitted visual observation. However, it remains unclear whether this mechanism is robust across laboratory settings. We aimed to replicate the findings by systematically investigating the role of visual observation in maternal behavior acquisition in pup-naive virgin female mice. Following the same protocol as previous studies, we compared retrieval performance under conditions that either allowed or prevented visual observation of maternal behavior. Contrary to previous findings reporting substantial differences between these conditions, virgin females readily acquired pup retrieval regardless of visual access to experienced mothers. We detected no barrier-related differences in retrieval success or in its trajectory across sessions. Barrier condition did affect the timing of successful retrievals: virgins in the opaque condition took longer to reach the pup on the first testing session only, whereas across all sessions they returned pups to the nest faster than virgins in the transparent condition, a direction not predicted by visual social transmission. These results question the replicability of visual transmission of maternal behavior in virgin female mice. Further discussion and examination is needed to determine whether visual observation plays a role in mouse alloparental behavior.

## Introduction

Animals possess a variety of innate behaviors that allow for their survival, and they also develop new behavioral repertoires through experience and exposure. An interesting behavior that falls into both categories is parental behavior, which animals express differently depending on sex and socio-sexual experience (Dulac et al., 2014; Kohl, 2020). In rodents, pup retrieval, the act of returning displaced pups to the nest, serves as a commonly used model for studying whether and how experience modulates innate behaviors. This behavior, consisting of locating pups astray from the nest, picking them up, and returning them, is particularly valuable because it is both stereotyped and measurable. Mothers promptly rescue stray pups back to the nest to avoid losing pups to death by hypothermia, and this behavior can also be expressed by virgin females, although at a smaller rate than mothers (Stolzenberg and Rissman, 2011 [rats], Marlin et al., 2015).

Two main hypotheses have been proposed to explain how pup retrieval emerges in alloparental subjects such as virgin females. The first posits that repeated exposure to pup-related sensory cues (odor, ultrasonic vocalizations, and tactile contact) is sufficient to sensitize naive animals and activate latent retrieval behaviors (Stern, 1997 [rats], Okabe et al., 2017; Schiavo et al., 2020). The second proposes that social transmission, specifically the visual observation of an experienced conspecific performing the behavior (Carcea et al., 2021), plays a critical role in behavioral acquisition. These hypotheses are not mutually exclusive, but their relative contributions remain unclear, and the conditions under which each mechanism is sufficient or necessary have not been established.

A recent study by Carcea et al., 2021 provided evidence in favor of the social transmission hypothesis, reporting that virgin females learned to retrieve pups when a transparent barrier allowed them to observe a demonstrating mother, whereas most failed when the barrier was opaque. This study makes a strong claim for the social transmission hypothesis in a highly ethologically-relevant behavior, and as such is worth validating.

In the present study, we tested whether visual access to maternal demonstrations was associated with pup-retrieval acquisition in virgin female mice, replicating the design of Carcea et al., 2021. Following the aforementioned protocol and employing completely pup-naive virgins weaned into a pup-free room, we compared the performance of virgin females in transparent versus opaque barrier conditions across multiple sessions. Virgins acquired retrieval behavior in both conditions regardless of visual access, and we detected no barrier-related difference in retrieval success or in its trajectory across sessions. Barrier condition did affect the timing of successful retrievals, in two ways. First, on the first testing session only, virgins in the opaque condition took longer than those in the transparent condition to reach the displaced pup. Second, across all sessions, virgins in the opaque condition returned pups to the nest faster. These findings are consistent with the pup-exposure hypothesis and question the robustness of a large social transmission effect under these experimental conditions.

## Results

### Naive females acquire pup retrieval behavior without a detectable effect of visual observation

The pup retrieval task consisted of two consecutive phases: a “demonstration” and a “testing” phase. In the demonstration phase, a mother and a virgin were placed on opposite sides of an arena divided by a barrier, and the mother retrieved displaced pups over 10 trials while the virgin remained on the other side. In the testing phase, the virgin was then tested for pup retrieval in the same manner (Fig. 1A). Each mother-virgin pair underwent four consecutive days of testing (d1–d4) after a prescreening session (d0) to assess baseline retrieval performance in each animal individually. To investigate whether visual observation influences pup retrieval acquisition, we assigned mother–virgin pairs to either a transparent (N = 14 pairs) or opaque barrier (N = 13 pairs) condition. In the transparent condition, virgins could observe maternal demonstrations, whereas in the opaque condition, visual observation was blocked (Fig. 1A). Both groups of virgins acquired retrieval behavior: 12 out of 14 virgins (85.7%) in the transparent group and 10 out of 13 virgins (76.9%) in the opaque group retrieved at least one pup in at least one session (Fig. 1B, C). The proportion of virgins retrieving did not differ between the transparent and opaque barrier conditions (two-sided Fisher’s exact test, *p* = .648).

**Figure 1:**
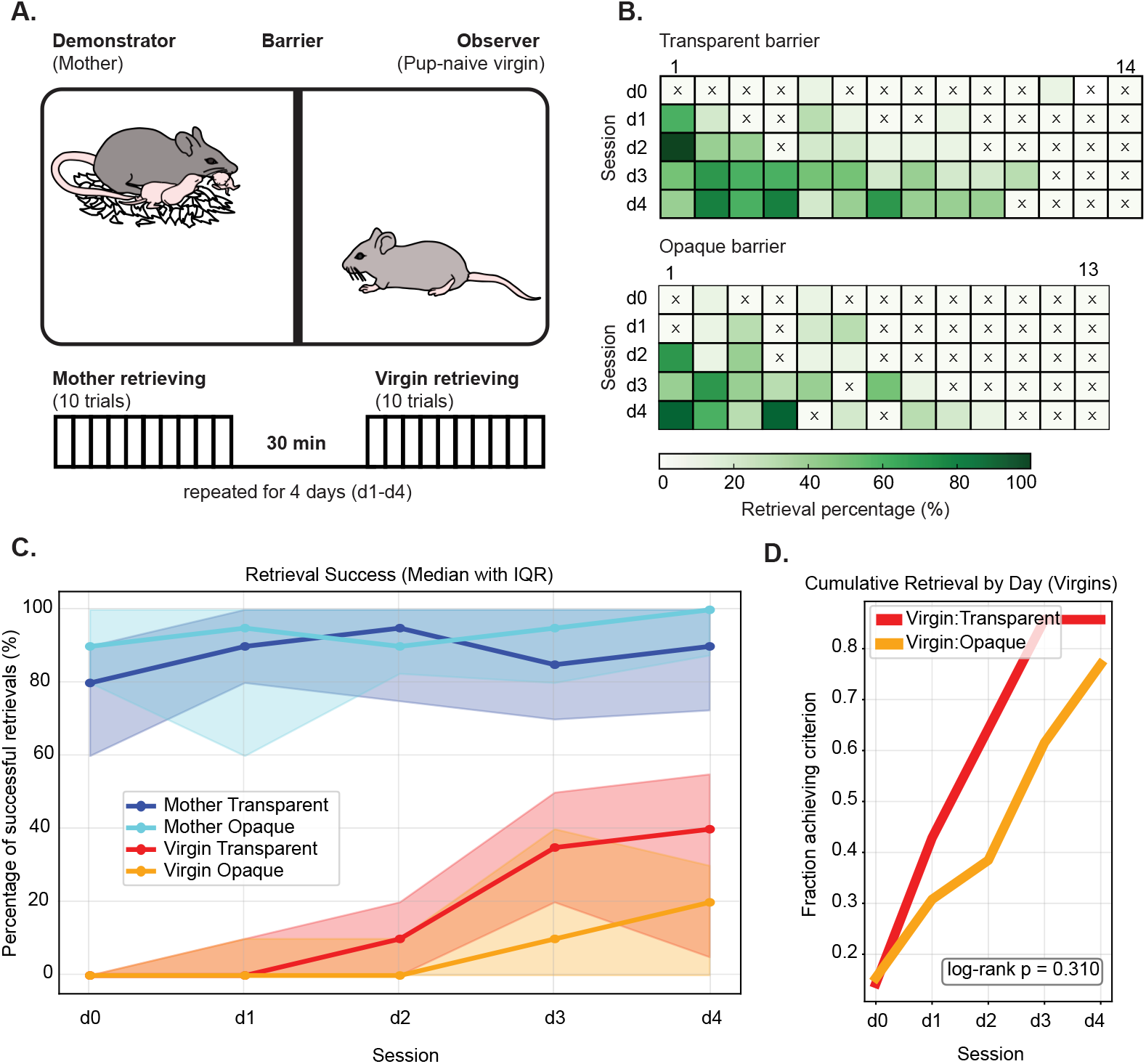
Retrieval success increases across sessions without a detectable barrier-dependent trajectory. A. Task schematic of pup retrieval behavior task. A mother demonstrates pup retrieval for 10 trials, while a virgin observes through a transparent or opaque barrier. After a 30-minute break, the virgin is then tested for pup retrieval. The whole paradigm is repeated for 4 days. **B**. Heatmaps show virgin retrieval performance in transparent condition (top plot, *N* = 14 animals) and opaque condition (bottom plot, *N* = 13 animals). Dark green color = 100% retrieval rate; white color with ‘X’ mark = 0% retrieval rate. **C**. Lineplots depict successful retrievals over days (median with IQR) for each experimental group (Mother: Transparent in blue, Mother: Opaque in cyan, Virgin: Transparent in red, Virgin: Opaque in orange). **D**. Cumulative proportion of virgins meeting the retrieval criterion: Virgins:Transparent (red) and Virgins:Opaque (orange). The log-rank test did not show any statistically significant difference between the two barrier conditions (log-rank *p* = 0.310).

Similarly, the onset of virgin retrieval did not differ based on barrier type (Fig. 1D, log-rank test, p=0.310). Four virgins (two per barrier group) retrieved one pup on d0; following the inclusion criterion of Carcea et al., 2021, virgins that retrieved more than one pup on d0 had been excluded (see Methods). To replicate the analysis approach of Carcea et al., 2021, we performed a two-way ANOVA on virgin retrieval success. For each virgin and session, we defined the success rate as the percentage of the 10 test trials in that session in which the virgin returned the displaced pup to the nest. The ANOVA included barrier type (transparent, opaque) and session (d0–d4) as factors. Session had a significant effect (*F* (4, 125) = 9.47, *p <* .001), whereas barrier type did not (*F* (1, 125) = 2.83, *p* = .095), and there was no significant barrier *×* session interaction (*F* (4, 125) = 0.34, *p* = .854; Extended Data Table 1). Thus, performance was associated with session, whereas this analysis detected neither an overall barrier effect nor a barrier-dependent session profile.

Despite similar overall rates, the two groups might differ in how retrieval was acquired: virgins in the opaque condition might learn more slowly, reflecting an effect of barrier condition on the rate of acquisition rather than on its endpoint. Such a difference would appear as a barrier *×* session interaction. Although we used the two-way ANOVA to replicate Carcea et al., 2021, this approach is not appropriate for these data. It treats each animal-session as an independent observation, although each virgin was tested in every session, so session is a repeated (within-subject) measure. It also treats success rates as continuous, normally distributed values, although they are counts of successes out of 10 trials.

Thus, to test whether barrier condition affected the acquisition trajectory, we fitted a binomial generalized linear mixed model (GLMM) with a logit link to the numbers of successful and unsuccessful trials for each animal-session. Barrier condition and session were included as fixed factors, together with their interaction, with session treated as a categorical variable so that acquisition was not constrained to a linear trajectory. Animal identity was included as a random intercept to account for the repeated measurements across sessions. In this model, the barrier *×* session interaction is the formal test of whether the across-session trajectory differs between barrier conditions.

Retrieval success differed significantly across sessions (main effect of session, Wald *χ*^2^(4) = 104.71, *p <* .0001; Table 1). Dunnett-adjusted comparisons with baseline (d0) showed significantly higher odds from d2 onward in the opaque condition and from d1 onward in the transparent condition (Table 2).

**Table 1:** Omnibus tests of barrier and session effects on retrieval success. Marginal type III Wald *χ*^2^ tests (via `emmeans::joint_tests`) from a binomial generalized linear mixed model (logit link) with barrier condition (transparent vs. opaque) and session (d0–d4) as fixed factors, together with their interaction, and animal identity as a random intercept (27 virgins total; 135 animal-session observations, 1,350 trials). The main effect of barrier compares transparent with opaque conditions, averaged equally across the five sessions; the main effect of session tests for differences among sessions, averaged equally across the two barrier conditions; and the barrier*×* session interaction tests whether the change across sessions differs between barrier conditions. Averages are taken over the model’s estimated marginal means on the log-odds scale. Treatment-coded coefficients, using opaque and d0 as reference levels, are reported in Extended Data Table 2.

| Effect | Wald $\chi^2$ | df | p |
| --- | --- | --- | --- |
| Barrier | 0.821 | 1 | .365 |
| Session | 104.708 | 4 | < .0001 |
| Barrier × Session | 0.776 | 4 | .942 |

**Table 2:**
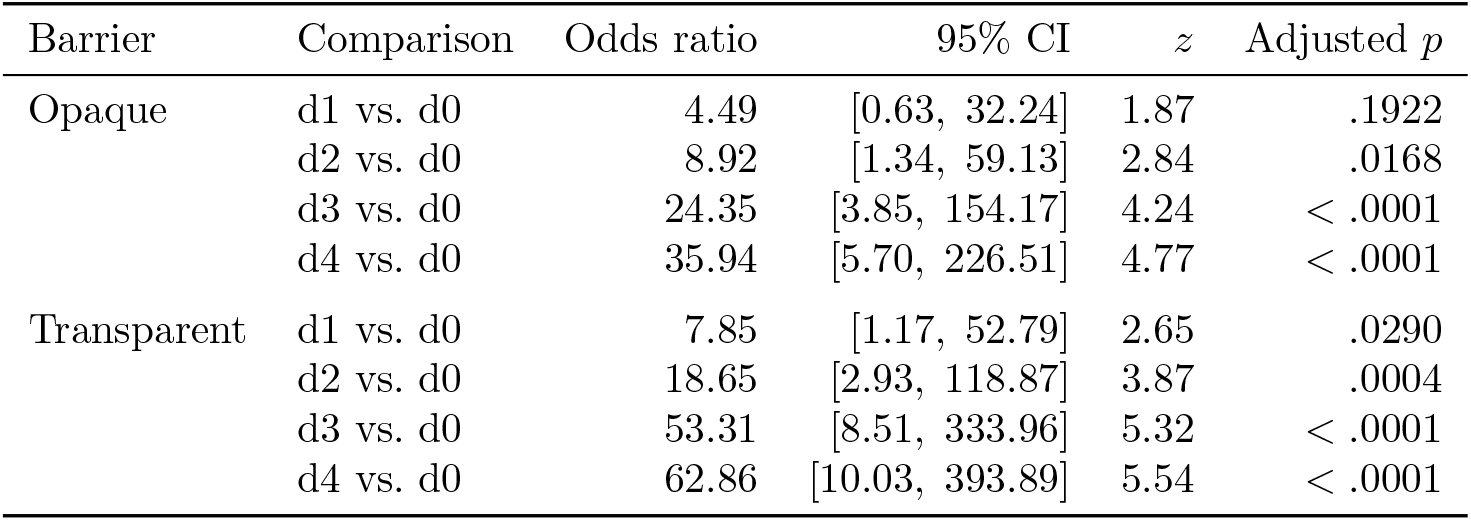
Primary comparisons of retrieval success across sessions. Estimated marginal odds ratios of successful retrieval in d1–d4 relative to d0 within each barrier condition. Confidence intervals and *p* values are Dunnett-adjusted for the four comparisons within each barrier. Odds ratios greater than 1 indicate higher odds of success than at d0 (which is the case for all of them). Tests were performed on the log-odds-ratio scale.

| Barrier | Comparison | Odds ratio | 95% CI | $z$ | Adjusted $p$ |
| --- | --- | --- | --- | --- | --- |
| Opaque | d1 vs. d0 | 4.49 | [0.63, 32.24] | 1.87 | .1922 |
|  | d2 vs. d0 | 8.92 | [1.34, 59.13] | 2.84 | .0168 |
|  | d3 vs. d0 | 24.35 | [3.85, 154.17] | 4.24 | < .0001 |
|  | d4 vs. d0 | 35.94 | [5.70, 226.51] | 4.77 | < .0001 |
| Transparent | d1 vs. d0 | 7.85 | [1.17, 52.79] | 2.65 | .0290 |
|  | d2 vs. d0 | 18.65 | [2.93, 118.87] | 3.87 | .0004 |
|  | d3 vs. d0 | 53.31 | [8.51, 333.96] | 5.32 | < .0001 |
|  | d4 vs. d0 | 62.86 | [10.03, 393.89] | 5.54 | < .0001 |

There was no significant main effect of barrier condition on retrieval success (*χ*^2^(1) = 0.82, *p* = .365), and the barrier *×* session interaction was not significant (*χ*^2^(4) = 0.78, *p* = .942; Table 1, full model coefficients in Extended Data Table 2). Holm-adjusted transparent-versus-opaque comparisons were nonsignificant at every session (all adjusted *p* = 1.000; Table 3). These comparisons were performed to mirror Carcea et al., 2021’s session-by-session analysis and are therefore reported regardless of the omnibus interaction result. Thus, retrieval success increased across sessions, but we did not detect a barrier-dependent acquisition trajectory. However, the broad confidence intervals indicate substantial uncertainty, and these nonsignificant results should not be interpreted as evidence of equivalence between conditions.

**Table 3:** Barrier comparisons of retrieval success within sessions. Odds ratios compare transparent with opaque barriers. Confidence intervals are Bonferroni-adjusted and *p* values are Holm-adjusted across the five session-specific comparisons. Odds ratios greater than 1 indicate higher odds of retrieval with the transparent barrier.

| Session | Odds ratio | 95% CI | $z$ | Holm-adjusted $p$ |
| --- | --- | --- | --- | --- |
| d0 | 1.08 | [0.05, 23.20] | 0.07 | 1.000 |
| d1 | 1.89 | [0.25, 14.57] | 0.80 | 1.000 |
| d2 | 2.26 | [0.34, 14.90] | 1.12 | 1.000 |
| d3 | 2.37 | [0.40, 14.03] | 1.25 | 1.000 |
| d4 | 1.89 | [0.33, 11.00] | 0.93 | 1.000 |

If visual observation contributed to the improvement in virgin performance across sessions, then virgins in the transparent condition should have performed better when they had observed a more proficient demonstrator. We therefore asked whether the quality of maternal demonstration predicted virgin performance. Mothers showed some variability in their retrieval performance over days (IQR shaded area on Fig. 1C). We therefore correlated session-by-session performance of mothers and virgins using Spearman rank correlations (Extended Data Fig. 1B). In the transparent-barrier condition, correlations ranged from *ρ* = *−*0.39 to *−*0.06; in the opaque-barrier condition, they ranged from *ρ* = *−*0.39 to 0.36. All correlations were non-significant after Benjamini–Hochberg correction across the eight session-specific tests (all false-discovery-rate-adjusted *p ≥* .498). These findings indicate that the quality of maternal demonstrations did not predict virgin performance.

Collectively, these results provide no evidence of a large benefit of visual social transmission under the conditions tested in Carcea et al., 2021. The consistently high number of retrievers in both barrier conditions, together with the absence of a detectable association between maternal and virgin performance, are consistent with pup exposure explaining retrieval acquisition.

### Barrier condition affects return latency throughout, and reach latency at first exposure

Having found no detectable effect of barrier condition on retrieval success, we next asked whether barrier transparency affected the timing of successful retrievals by virgins. We divided retrieval into two phases: “reach pup latency,” from pup displacement until the virgin picked up the pup, and “return pup latency,” from pick-up until the pup was returned to the nest (Fig. 2). Because d0 preceded the demonstration manipulation and latency was measurable for only four animals at d0 (i.e., successfully retrieved on at least one trial), the primary latency analyses included experimental sessions d1–d4 only. For each session of each animal, we calculated the median latency over successful trials.

**Figure 2:**
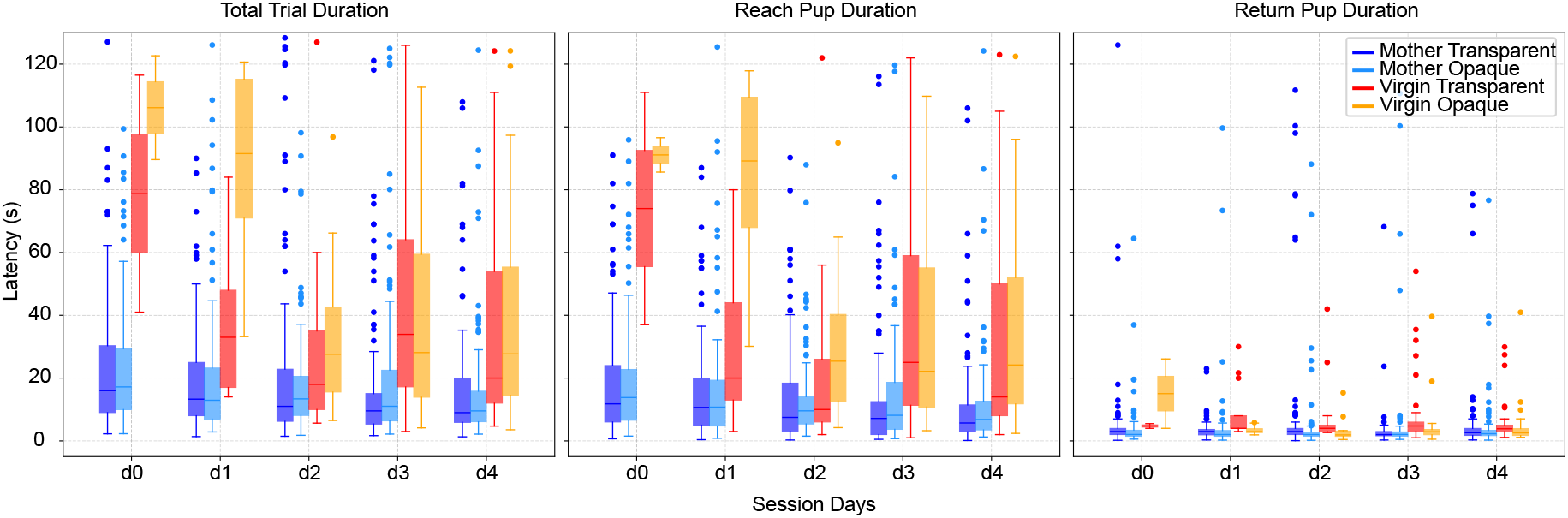
Retrieval-latency distributions across experimental groups. Boxplots show distributions of individual successful-trial latencies; boxes indicate medians and interquartile ranges. Latency and IQR for **Left**. total trial duration, **Middle**. “reach pup latency,” and **Right**. “return pup latency.” Inferential comparisons of opaque and transparent virgin groups during d1–d4 are reported in Table 4.

**Table 4:** Omnibus tests of barrier and session effects on reach and return latency in virgin mice. Type III *F* -tests (Satterthwaite denominator degrees of freedom) from linear mixed-effects models of animal-session median latency, with barrier and session as fixed factors, together with their interaction, and animal as a random intercept. Reach latency was modeled on its raw scale and return latency on the log scale, based on residual diagnostics (see Methods). Analyses included experimental sessions d1–d4 and 58 animal-session observations from 21 virgins for each outcome, and are conditional on at least one successful retrieval in an animal-session. Main effects are averaged equally across the levels of the other factor. *p* values for each term are Holm-adjusted across the two outcomes. Session-specific barrier contrasts for reach latency are reported in Extended Data Table 3.

| Outcome | Effect | $F$ | $df$ | $p$ | Holm $p$ |
| --- | --- | --- | --- | --- | --- |
| Reach (raw scale) | Barrier | 8.13 | 1, 19.0 | .010 | .010 |
|  | Session | 7.57 | 3, 33.6 | < .001 | .001 |
| | Barrier × Session | 7.60 | 3, 33.6 | < .001 | .001 |
| Return (log scale) | Barrier | 11.16 | 1, 13.3 | .005 | .010 |
|  | Session | 0.41 | 3, 36.6 | .748 | .748 |
| | Barrier × Session | 1.25 | 3, 36.6 | .305 | .305 |

To test whether barrier condition affected the timing of successful retrievals, and whether any such effect changed across sessions, we fitted a linear mixed-effects model for each outcome (animal-session median reach latency and median return latency). Each model included barrier and session as fixed factors, together with their interaction, and animal identity as a random intercept. Model scale was selected independently for each outcome based on residual diagnostics: reach latency was modeled on its raw scale, and return latency on the log scale (see Methods). For each model, we first tested the main effects of barrier and session and the barrier *×* session interaction (Table 4). Because we tested the same terms in two models (reach and return), we corrected for multiple comparisons by Holm-adjusting the *p* values for each term across the two outcomes. Follow-up comparisons depended on the interaction: where it was significant, we compared the barrier conditions within each session; where it was not, we estimated a single session-averaged barrier effect. For reach latency, the barrier *×* session interaction was significant (*F* (3, 33.6) = 7.60, *p*_Holm_ = .001; Table 4), indicating that the barrier effect on reach latency changed across sessions. The main effects of session (*F* (3, 33.6) = 7.57, *p*_Holm_ = .001) and barrier (*F* (1, 19.0) = 8.13, *p*_Holm_ = .010) were also significant. Because of the interaction, however, the barrier main effect (an average over sessions) does not describe a consistent difference between conditions, so we compared the barrier conditions within each session. Opaque-condition virgins took markedly longer to reach the pup than transparent-condition virgins at d1 (86.7 vs. 29.0 s; *t*(47.6) = 4.48, *p*_Holm_ *<* .001), whereas the difference was not significant at d2, d3, or d4 (all *p*_Holm_ *≥* .119; Extended Data Table 3). Thus, the barrier effect on reach latency was confined to the first testing session rather than being a sustained effect on approach behavior.

For return latency, the main effect of barrier was significant (*F* (1, 13.3) = 11.16, *p*_Holm_ = .010), whereas neither the main effect of session (*F* (3, 36.6) = 0.41, *p*_Holm_ = .748) nor the barrier *×* session interaction (*F* (3, 36.6) = 1.25, *p*_Holm_ = .305) was significant (Table 4). Because the barrier effect did not detectably differ across sessions, we estimated it from a model without the interaction term. Return latency was shorter in the opaque condition than in the transparent condition (session-adjusted geometric means 3.09 vs. 4.87 s; opaque-to-transparent ratio 0.634, 95% CI [0.469, 0.858], *t*(16.4) = *−*3.19, *p*_Holm_ = .011). A parametric bootstrap likelihood-ratio test for the barrier term in this model (5,000 simulations under the model without barrier) confirmed this effect (LRT = 8.84, *p*_Holm_ = .012). To confirm that summarising each animal-session by its median latency does not introduce distortions to results, a trial-level sensitivity analysis retained all successful trials per outcome and included random intercepts for animal and animal-session, with reach modeled on the raw scale and return on the log scale to match the primary models (see Methods and Extended Data Table 4). The trial-level results matched the animal-session median analysis: the barrier *×* session interaction was significant for reach (*p*_Holm_ = .034) but not for return (*p*_Holm_ = .548), and the session-averaged barrier effect on return latency was significant (*p*_Holm_ = .002). These results corroborate a session-dependent barrier effect on reach latency and a stable barrier effect on return latency.

## Discussion

Our study investigated whether visual observation of maternal pup retrieval behavior promotes acquisition in completely pup-naive virgin female mice, using the same protocol as Carcea et al., 2021. Virgin females demonstrated robust pup retrieval regardless of barrier transparency: 85.7% of virgins retrieved in the transparent condition and 76.9% in the opaque condition, with no significant difference between the groups (Fig. 1). We followed the full experimental design of Carcea et al., 2021, including their criterion for excluding virgins that retrieved more than one pup at prescreening (d0). Our results therefore cannot be explained by the inclusion of virgins that already retrieved readily before the experiment. Separately, a larger fraction of our prescreened virgins retrieved at least one pup at d0 (approximately 20%) than in Carcea et al., 2021 (less than 5%).

Our results are consistent with established pup sensitization in virgins (Okabe et al., 2013; Rosenblatt, 1967; Stern, 1997; Stolzenberg and Rissman, 2011), but diverge from Carcea et al., 2021, in which 63% of virgins in the transparent condition and 11% in the opaque condition acquired retrieval (compared with 85.7% and 76.9%, respectively, here). In our data, descriptive trends slightly favored the transparent barrier condition (e.g., d3 having 11/14 retrievers in transparent barrier and 7/13 in opaque), yet the retrieval-success model showed no significant barrier effect or barrier *×* session interaction. Taken together, these results suggest that visual social transmission of pup retrieval may be less robust than previously reported, and that conclusions drawn from small-sample studies should be interpreted with caution.

Interpreting these findings in the context of existing literature requires attention to substantial methodological heterogeneity across studies. Existing protocols differ markedly in the extent to which virgins are exposed to pup-related cues before testing. Some studies explicitly report using virgins raised completely isolated from pups (Stolzenberg and Rissman, 2011, Carcea et al., 2021; Schiavo et al., 2020), whereas others provide no clear information about such isolation (Rosenblatt, 1967 [rats], Glat et al., 2022; Marlin et al., 2015; Muzerelle et al., 2021; Okabe et al., 2013, 2017). Other studies include virgins that were co-housed with mothers and pups (Stern, 1997); Carcea et al., 2021 used both pup-isolated and co-housed virgins. This heterogeneity complicates direct comparisons across studies and raises the possibility that prior exposure to pup cues - even indirect exposure via olfactory or auditory signals - may shape the likelihood and time course of pup retrieval in sexually-naive animals. Our subjects (pup-naive virgins), weaned into a pup-free room, thus strengthen the evidence that pup cue exposure alone can suffice to trigger retrieval behavior in alloparental subjects, while also highlighting the sensitivity of this task to housing and procedural factors.

The latency analyses add an important qualification to the retrieval-success results: barrier condition was associated with the timing of successful retrievals, but its influence differed between the two phases of the behavior. For reach latency, opaque-condition virgins were markedly slower to reach the pup at d1, but the difference was not significant from d2 onward. This pattern is consistent with a transient effect of barrier condition on approach behavior during the first testing session, plausibly reflecting differences in novelty or arousal associated with the two barrier types, rather than a sustained influence on reach behavior. In contrast, return latency was shorter in the opaque condition throughout d1–d4, with no evidence of a barrier *×* session interaction.

The shorter return latency in the opaque condition was not predicted by the visual social-transmission hypothesis and should not be interpreted as evidence that visual information impairs retrieval. Barrier transparency may alter attention, arousal, or other features of the testing context, but the present behavioral measurements cannot distinguish among these possibilities. Future work would be required to establish whether this timing effect is reproducible and which sensory or contextual processes produce it.

Several features of the data limit the latency analyses. Latency was observable only for successful retrievals, so the latency models describe the subset of virgins and animal-sessions in which retrieval occurred; latency cannot be estimated for failed animal-session trials. In addition, the number of successful trials contributing to each animal-session median varied across animals and sessions. Latencies were also recorded at a coarser resolution in the transparent than in the opaque condition, so a systematic difference in manual scoring between conditions cannot be fully excluded. The trial-level sensitivity models yielded similar effect estimates but had singular random-effect structures. More broadly, our conclusions rest on behavioral measures alone, without direct assessment of neural activity, hormonal states, or genetic factors, and larger samples with measurements capable of accommodating unsuccessful trials will be needed to establish the robustness and mechanism of the observed return-latency difference.

The similar acquisition rates in the transparent and opaque conditions suggest that direct exposure to pups during testing, rather than visual observation of a demonstrating mother, was sufficient for virgins to acquire retrieval. Future experiments should identify which pup-derived cues drive this acquisition. Pup odors and ultrasonic vocalizations can trigger maternal responses in virgin females (Marlin et al., 2015; Okabe et al., 2013; Schiavo et al., 2020), and tactile contact with pups may also contribute. Note that the opaque barrier blocked visual access to the mother, but not necessarily the sounds and odors of the mother and litter during demonstration. Our design therefore cannot exclude social transmission through non-visual channels; testing this possibility would require a demonstration phase that also blocks auditory and olfactory cues.

## Methods

### Animals

All of the procedures and experiments were approved by the Cantonal Veterinary Office Zurich, Switzerland. All experiments used female C57Bl/6J mice (Jackson Lab) aged between 7 and 22 weeks at the start of the behavioral experiment. Animals were group-housed (around 2–4 same-sex litter-mates) in individually ventilated cages (IVC) in a 12 h light/dark cycle room, and we provided them with food and water *ad libitum*. The subjects fell into two groups: mothers (either primiparous or multiparous at the start of the experiment) and sexually- and pup-naive females (virgins), which were housed separately. Virgins were bred in-house, and we transferred them to a different pup- and breeding-free room upon weaning to avoid exposure to other pups and breedings. Virgins remained male-naive throughout the experiments. Mothers and virgins were prescreened (“day zero”, d0) for retrieval and pup mauling before the behavioral task began (see Behavioral task). Following Carcea et al., 2021, virgins that retrieved more than one pup at d0 were excluded from the study.

### Behavioral task

The established pup retrieval task (Carcea et al., 2021) consists of two consecutive stages, “demonstration” and “testing”, which we repeated over 4 days after prescreening (postnatal days 1–4 of mother’s pups). During prescreening (d0), each mother and each virgin was tested individually for pup retrieval in 10 trials, following the testing procedure described below but without a preceding demonstration; animals were also monitored for pup mauling. During the ‘demonstration’ phase, the mother and the virgin were placed on opposite sides of a conventional 1800 cm^2^ home cage (588 mm *×* 194 mm *×* 395 mm, NexGen IVC for rats 1800, Allentown), and an opaque or transparent barrier separated them at the midline. We gave the animals 20 minutes to acclimatize before each demonstration session began. The mother’s entire litter was grouped in a corner of the arena on the mother’s side, and we covered them with nesting material. After an additional 1 – 2 minutes of acclimatization, we removed one pup from the nest and pseudorandomly placed it in either of the 3 other corners of the mother’s cage partition, and we gave the mother a maximum of 2 minutes and 5 seconds to retrieve the displaced pup back to the nest. After 10 trials, the animals returned to their home cage, and a 30-minute break followed. We changed the used cage into a new, clean cage, and we placed the virgin on one of the partitions of the arena. After 20 minutes of acclimatization, the “testing” phase began, and we put the entire litter back into the corner of the testing arena, covering them with nesting material. After 1 – 2 minutes of acclimatization, we tested the virgin for pup retrieval in 10 trials, as we described above. At the end of the experiment, all the animals returned to their home cages. We applied two different conditions of this task: transparent barrier, where the barrier dividing the cage was transparent (and therefore the virgins could observe the mother) and an opaque condition, which consequently impeded the virgin from observing the mother and litter.

### Behavioral analysis

A top view B/W CMOS camera (60516, Stoelting Europe) integrated with an ANY-maze system (Stoelting Europe) recorded all mouse behavior. The following parameters were quantified for analysis: **Reach pup latency**: the time to approach the pup ending in a pick-up, from pup displacement from the nest to the pick-up. **Return pup latency**: the time to retrieve the pup to the nest, from pick-up to return to the nest. **Total latency**: the whole trial duration, which is the sum of reach pup latency and return pup latency. **Retrieval success**: we categorized a trial as ‘successful’ if the female (mother/virgin) brought the pup back to the nest within the time allotted (maximum 2 minutes and 5 seconds), and ‘failed’ otherwise (the experimenter then returned the pup to the nest at the end of the allotted time). In six of 1,354 successful trials (two of them virgin trials), the pup was returned to the nest slightly after the limit (within 2 minutes and 9 seconds); these trials were counted as successful.

### Statistical analysis

Acquisition of pup retrieval by virgins was compared using a two-sided Fisher’s exact test on the number of retrievers in each barrier condition. The onset of retrieval was compared using a log-rank test on the Kaplan–Meier curves for each barrier condition, with onset defined as the first session (d0–d4) in which a virgin retrieved at least one of the 10 pups; virgins that never met this criterion were censored at d4. To mirror the analysis in Carcea et al., 2021, we also conducted a two-way ANOVA on virgin success rates, with barrier type (transparent, opaque) and session (d0–d4) as factors. The success rate of each virgin in each session was defined as the percentage of the 10 test trials in that session in which the virgin returned the displaced pup to the nest. The ANOVA used Type I (sequential) sums of squares, which equal Type II sums of squares here because every virgin was tested in every session.

We fit the mixed-effects models in R using the `lme4` package and used the `emmeans` package for marginal omnibus tests. For the linear mixed-effects models, coefficient-level significance tests were obtained using `lmerTest`. Estimated marginal means and contrasts were calculated using `emmeans`, and simulation-based residual diagnostics were performed using `DHARMa`. All remaining analyses, including the two-way ANOVA and Spearman rank correlations described below, were performed in Python.

To assess whether barrier condition influenced retrieval success across sessions, we fitted a binomial generalized linear mixed-effects model (GLMM) with a logit link using `glmer` from `lme4`. For each animal and session, the numbers of successful and unsuccessful retrievals across 10 trials were entered as a grouped response. Barrier condition (transparent or opaque) and session were included as fixed factors, together with their interaction. Animal identity was included as a random intercept to account for repeated measurements. Session was treated as a categorical factor so that acquisition was not constrained to follow a linear trajectory. The model was fitted by maximum likelihood using the Laplace approximation.

The regular binomial model showed extra-binomial variation (Pearson dispersion ratio = 1.86, *p <* .001; `DHARMa` dispersion test, *p* = .048) and a mild gradient convergence warning. The Q–Q and residual-versus-fitted plots were therefore inspected for unusually large residuals and systematic patterns. Because Pearson residuals from a binomial GLMM are discrete and are not expected to be normally distributed, the Q–Q plot was treated as an outlier screen rather than a formal normality test. These diagnostic limitations were considered when interpreting the model estimates.

Barrier and session were coded using treatment contrasts, with opaque and d0 as the reference levels. Fixed effects were evaluated using marginal omnibus tests implemented with `emmeans::joint_tests` (Table 1). These tests are computed from the estimated marginal means (log-odds scale) and therefore do not depend on the choice of reference levels. The main effect of barrier compares transparent with opaque conditions, averaged equally across sessions; the main effect of session tests for differences among sessions, averaged equally across barrier conditions; and the interaction tests whether the change across sessions differs between barrier conditions. Treatment-coded coefficients from the same model, using opaque and d0 as reference levels, are reported in Extended Data Table 2. For planned comparisons, sessions d1–d4 were compared with d0 separately within each barrier, with Dunnett adjustment across the four comparisons in each barrier (Table 2). Transparent and opaque barriers were compared within each session, with Holm adjustment across the five tests (Table 3).

We analyzed reach and return latencies among virgin animals during experimental sessions d1–d4. Session d0 was excluded because it preceded the demonstration manipulation and latency was measurable for only four successful animals at baseline. For each animal, session, and outcome, we calculated the median latency across successful trials; animal-sessions without a successful latency were absent.

Separate linear mixed-effects models were fitted for reach and return by restricted maximum likelihood (REML) with the `lmer` function (`lme4` and `lmerTest`). Model scale was selected independently for each outcome based on residual diagnostics on the d1–d4 subset. Reach latency was modeled on its raw (untransformed) scale, which showed acceptable residual symmetry and no systematic residual-versus-fitted pattern; log-transformation instead introduced a pronounced negative-tail residual. Return latency showed right-skew and heteroscedasticity on the raw scale; log-transformation substantially improved residual behavior and was used for this outcome. The model for each outcome was `[reach or log(return)]` *∼* `barrier * session + (1` | `animal)`, with session treated categorically and barrier and session coded using treatment contrasts (opaque and d1 as reference levels). The main effects of barrier and session and the barrier *×* session interaction were tested with Type III *F* -tests with Satterthwaite denominator degrees of freedom (`lmerTest`), which do not depend on the choice of reference levels. For each term, *p* values were Holm-adjusted across the two outcomes. Where the interaction was significant (reach), barrier contrasts were computed within each session from the interaction model using `emmeans` with Kenward-Roger degrees of freedom and Holm-adjusted across the four sessions. Where the interaction was not significant (return), the session-averaged barrier effect was estimated from the corresponding additive model (`barrier + session + (1` | `animal)`) using `emmeans` with Kenward-Roger degrees of freedom, with Holm adjustment across the two outcomes.

As a further check on the session-averaged barrier effect, we conducted a parametric bootstrap likelihood-ratio test for the barrier term in each additive model, comparing it with a reduced model omitting barrier (5,000 simulations under the reduced model, using `PBmodcomp` from `pbkrtest` with models fitted by maximum likelihood and a fixed random seed); Holm correction was applied across the two outcomes. As a trial-level sensitivity analysis, we retained each successful reach or return trial and fitted models with barrier and session as fixed effects and random intercepts for animal and animal-session. Secondary models added the barrier *×* session interaction. Holm correction was again applied across reach and return barrier tests. The trial-level models produced singular fits because the animal-session variance was estimated near zero for both outcomes and the animal variance was estimated at zero for return latency; they were therefore used to assess the stability of effect estimates rather than as the primary inferential models (Extended Data Table 4). Finally, we examined the relationship between maternal and virgin performance over days using session-specific Spearman rank correlations (Python) to test whether maternal success predicted virgin acquisition within pairs. The resulting eight *p* values (four sessions in each barrier condition) were adjusted together using the Benjamini–Hochberg false-discovery-rate procedure.

## Author contributions

All authors contributed to the study conception and design. Material preparation, data collection, and analysis were performed by E.S.M. The first draft of the manuscript was written by E.S.M. and P.V.A. and all authors commented on previous versions of the manuscript. All authors read and approved the final manuscript.

## Acknowledgments

The authors thank the UZH LASC team for animal husbandry and Krzysztof Cybulski for statistics assistance. This work was supported by the Swiss National Science Foundation (B.F.G. CRSII5-173721 and 315230 189251, P.V.A. 182539 funding), ETH project funding (B.F.G. ETH-20 19-01), and a Forschungskredit from the University of Zurich (P.V.A. FK-24-122)

## Extended Data

**Extended Data Fig. 1.**
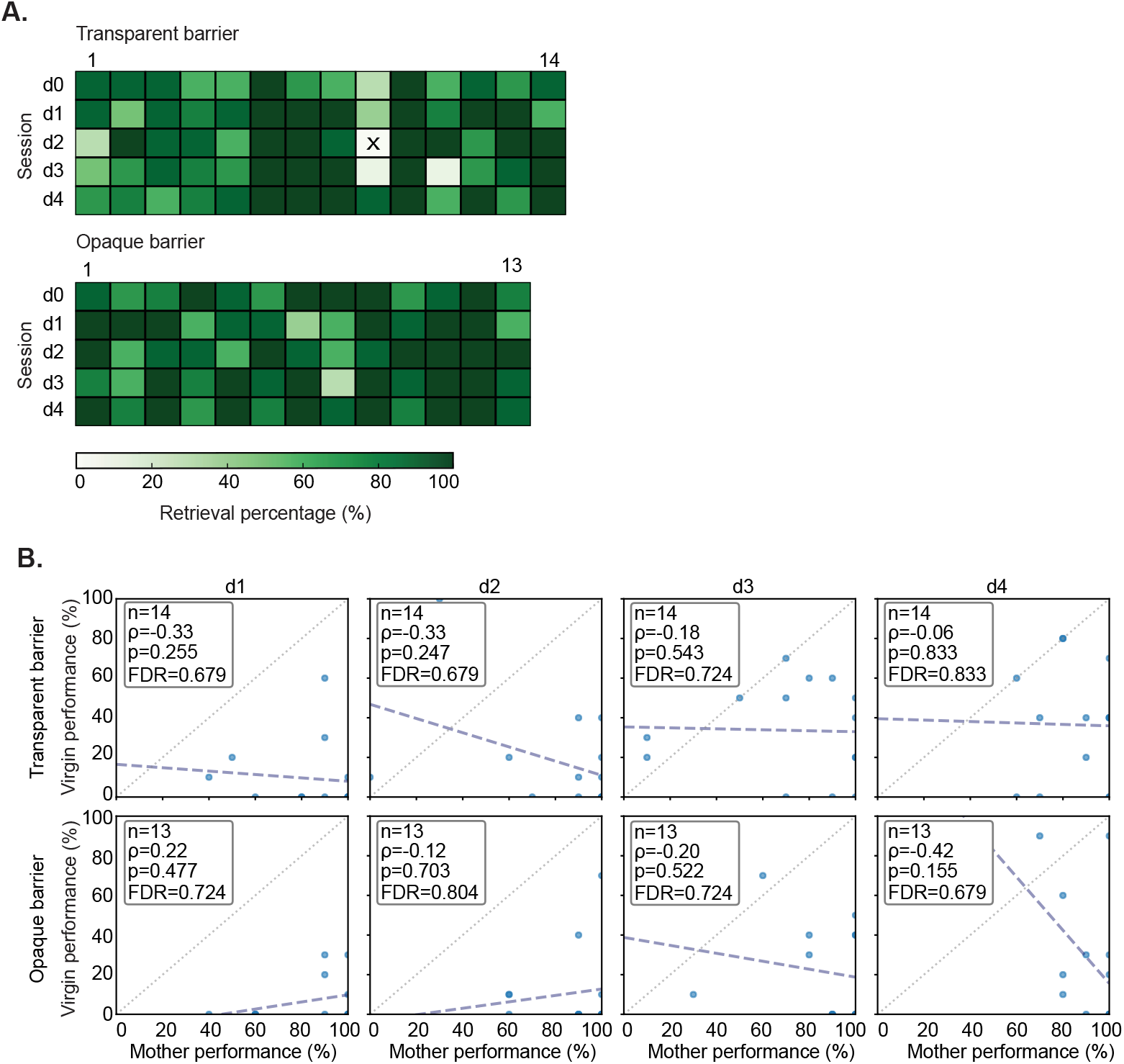
Maternal retrieval performance and mother–virgin performance correlations across barrier conditions. A. Heatmaps show maternal retrieval performance in transparent condition (top plot, *N* = 14 animals) and opaque condition (bottom plot, *N* = 13 animals). Dark green color = 100% retrieval rate; white color with ‘X’ mark = 0% retrieval rate. **B**. Session-by-session Spearman correlations between maternal and virgin performance. All eight correlations were non-significant after Benjamini–Hochberg correction.

