## Extended data and tables for "Pup Exposure Is Sufficient for Activating Pup Retrieval Behavior in Pup-Naive Virgin Mice"

| | DF | SS | MS | F | $p$ |
|---|---|---|---|---|---|
| Barrier | 1 | 1350.33 | 1350.33 | 2.83 | .095 |
| Session | 4 | 18051.85 | 4512.96 | 9.47 | $< .001$ |
| Barrier × Session | 4 | 639.58 | 159.89 | 0.34 | .854 |
| Residual | 125 | 59558.24 | 476.47 |  |  |

Extended Data Table 1 **Standard two-way ANOVA of virgin retrieval success**, included to replicate the analysis of Carcea et al., 2021. The outcome was the success rate of each virgin in each session (percentage of the 10 test trials with a successful retrieval; 27 virgins × 5 sessions = 135 observations), with barrier type and session (d0–d4) as factors. Session had a significant effect on retrieval success, whereas neither barrier type nor the barrier × session interaction was significant. This analysis treats the repeated sessions of each virgin as independent observations; the binomial GLMM (Table 1) accounts for them. DF: degrees of freedom, SS: sum of squares, MS: mean square, F: F-statistic.

| Model-fit statistic | Value |
|---|---|
| Observations | 135 |
| Animals | 27 |
| AIC | 433.2 |
| BIC | 465.1 |
| Log likelihood | −205.6 |
| Deviance | 411.2 |

| Group | Term | Variance | SD |
|---|---|---|---|
| Animal (pair_id) | Intercept | 2.400 | 1.549 |

| Fixed-effect coefficient | Estimate | SE | $z$ | $p$ |
|---|---|---|---|---|
| Intercept (opaque, d0) | −5.095 | 0.852 | −5.98 | < .001 |
| Transparent vs. opaque at d0 | 0.079 | 1.190 | 0.07 | .947 |
| d1 vs. d0 in opaque | 1.503 | 0.803 | 1.87 | .061 |
| d2 vs. d0 in opaque | 2.188 | 0.771 | 2.84 | .005 |
| d3 vs. d0 in opaque | 3.192 | 0.752 | 4.24 | < .001 |
| d4 vs. d0 in opaque | 3.582 | 0.750 | 4.77 | < .001 |
| Transparent × d1 (vs. opaque × d0) | 0.558 | 1.118 | 0.50 | .618 |
| Transparent × d2 (vs. opaque × d0) | 0.738 | 1.079 | 0.68 | .494 |
| Transparent × d3 (vs. opaque × d0) | 0.784 | 1.061 | 0.74 | .460 |
| Transparent × d4 (vs. opaque × d0) | 0.559 | 1.059 | 0.53 | .598 |

Extended Data Table 2 **Binomial generalized linear mixed-model summary for retrieval success.** The model used a logit link and Laplace approximation: `cbind(v_successes, v_failures)` ∼ `barrier` × `session + (1 | pair_id)`. Treatment contrasts used opaque and d0 as reference levels, and coefficients are on the log-odds scale. With this coding, the barrier coefficient is the transparent–opaque difference at d0 only, and the session coefficients are differences from d0 in the opaque condition only. Each interaction coefficient is the difference between barrier conditions in the change from d0 to the given session; for example, Transparent × d1 = $(\text{d1} - \text{d0})_{\text{transparent}} - (\text{d1} - \text{d0})_{\text{opaque}}$. None of these coefficients is a main effect; main effects are tested in Table 1. The random intercept is for each virgin (`pair_id` identifies the mother–virgin pair, which contains one virgin). Multiplicity-adjusted estimated-marginal-mean comparisons are reported in Tables 2 and 3. The initial fit produced a mild gradient convergence warning, and diagnostics indicated extra-binomial variation (Pearson dispersion ratio = 1.86, $p < .001$; `DHARMa` dispersion test, $p = .048$); inferential results should therefore be interpreted cautiously.

| Session | Opaque − Transparent (s) | $t$ | $df$ | Holm-adjusted $p$ |
|---|---|---|---|---|
| d1 | 57.78 | 4.48 | 47.6 | < .001 |
| d2 | 21.17 | 1.83 | 42.1 | .149 |
| d3 | −1.55 | −0.15 | 34.5 | .882 |
| d4 | 21.83 | 2.14 | 33.7 | .119 |

Extended Data Table 3 **Session-specific barrier contrasts for reach latency.** Estimated marginal differences (opaque minus transparent) in median reach latency, from a linear mixed-effects model of raw-scale reach latency with barrier and session as fixed factors, together with their interaction, and animal as a random intercept (sessions d1–d4). $p$ values are Holm-adjusted across the four session-specific comparisons (Kenward-Roger degrees of freedom). The barrier difference was significant only at d1.

| Outcome | Trials | Animals | Effect | 95% CI | Holm $p$ | Interaction Holm $p$ |
|---|---|---|---|---|---|---|
| Reach (s, raw scale) | 221 | 21 | 11.8 s longer, opaque | [−4.58, 28.15] | .147 | .034 |
| Return (ratio, log scale) | 221 | 21 | ratio 0.623 | [0.492, 0.789] | .002 | .548 |

Extended Data Table 4 **Trial-level sensitivity analysis of barrier effects on virgin retrieval latency.** Separate linear mixed-effects models retained every successful reach or return trial from experimental sessions d1–d4, with reach modeled on the raw scale and return on the log scale to match the primary animal-session median models. Models included barrier and session as fixed effects and random intercepts for animal and animal-session. For reach, the effect is the opaque-minus-transparent difference in seconds; for return, it is the opaque-to-transparent ratio, back-transformed from the log scale. Confidence intervals and $p$ values are from estimated marginal contrasts, with Holm correction across the two primary outcomes. The final column reports Holm-adjusted secondary barrier × session tests. The animal-session variance was estimated near zero for both outcomes, and the animal variance was estimated at zero for return latency, producing singular fits. Accordingly, these models are treated as sensitivity analyses. Effect estimates and significance patterns closely matched the primary animal-session median analysis for both outcomes, including a significant reach barrier × session interaction consistent with the primary model.
